# Machines learn ecological networks: automated discovery of ecological networks based on empirical data

**DOI:** 10.1101/2023.03.04.527358

**Authors:** Hongseok Ko, Ahyoung Amy Kim, Hao Helen Zhang

## Abstract

Constructing ecological networks is known to be important and challenging in community ecology. In particular, to construct the holistic structure of ecological networks, identifying species interaction is essential but often costly and impalpable. Recent studies providing major challenges in assembling ecological networks have highlighted the need of new and more powerful approaches to reconstruct biological networks, including species interaction networks. In literature, there are no promising verifications in using machine leaning (ML) approaches to reconstruct ecological networks. In this work, we develop and employ a variety of ML methods, including penalized regression and graphical tools, to reconstruct ecological networks. For evaluation, we apply the methods to empirical time series data sets of 20 species abundances collected at Lake Constance in central Europe. We use resampled data to identify highly-ranked interactions among species and measure their consistency across 7 ML methods and 5,000 learning processes. We show that the best precision, recall, and F1 score were 0.48, 0.97, and 0.64, respectively, among all penalized regression methods under comparison. In summary, our study shows that machine learning methods offer promising data-driven and automated tools for reconstructing ecological networks and discovering underlying biological interactions among species.

## 1. Introduction

In community ecology, understanding species interactions is critical to construct structure and dynamics of community network. Usually myriad of interactions among species constituting a tangled bank are not yet fully charted. Generally speaking, there are two major types of interactions among species: trophic and non-trophic interactions. Trophic interactions describe who eats whom relationships between species, while non-trophic interactions cover any non-feeding relationship among species. Non-trophic interactions can be categorized by their interaction type and strength, for example, hummingbirds pollinating flowers, plants or microorganisms releasing allelopathic chemicals into the environment causing positive or negative interactions between them, sea anemones competing for space in tide pools, etc. It is believed that there are more abundances and types of non-trophic interactions in the real world compared to those of trophic interactions (Ellison, 2019; Kéfi et al., 2012).

To identify species interactions and study population dynamics, a large amount of ecological data has been collected in literature for many years, at a high cost of time-consuming and expensive human effort. However, there are several major challenges in constructing ecological networks: (i) the ecosystem is typically very large, but only limited empirical data are available, making it impracticable to assemble the complete community networks; (ii) the ecosystem is too complex to enable researchers to build accurate and highly interpretable models for estimating and predicting the underlying network structure; (iii) non-trophic interdependent relationships are much harder to observe or measure than trophic ones (Ellison, 2019). Furthermore, due to responding to changes in biotic and/or abiotic factors, these interactions can be altered, emerge, or vanish, and learning dynamic networks is even more difficult with a limited number of observations.

To resolve the above difficulties, one common strategy used in literature is to focus on only a small subset of species interactions at a time. Though it provides insight on local or partial structures of the network, it can not learn the entire network structure. In order to estimate large-scale assemblages of species interactions and discover potential interactions that have not been discovered, there is a need for an innovative way of constructing the community network structure from empirical observations. Modern machine learning methods provide powerful tools to learn complex networks from large volumes of data and achieve high prediction accuracy and efficiency (Sachs et al. 2005; Marbach et al. 2012; Camacho et al. 2018). In literature, a variety of machine learning methods have been developed to acquire information about the links (i.e., interactions) between nodes (e.g., species, genes, proteins) and reconstruct networks from high-dimensional data, including high-dimensional penalized linear regression (Kolar & Xing, 2012; Meinshausen & Bühlmann, 2006; Yang & Peng, 2019) and graphical models (Fan et al., 2009; Friedman et al., 2008; Li & Gui, 2006).

However, among existing machine learning algorithms to infer species interactions based on empirical data, very few attempts were made to exploit presence-absence data (Faisal et al., 2010; Sander et al., 2017) or time series data of species abundance (Faisal et al. 2010; Bucci et. al. 2016). In this study, we will study a set of machine learning methods for automatically learning species interactions and reconstructing inclusive ecological networks based on empirical presence-absence data. In particular, we consider five state-of-the-art penalized regression methods, including LASSO (Tibshirani, 1996) and SCAD (Fan & Li, 2001), and two graphical models to reconstruct and visualize ecological networks. There are a few advantages with the new methods. First, they can automate certain tasks, such as variable selection and covariance selection, to detect edges connecting nodes and communities, and are therefore computationally scalable for large networks. Second, instead of focusing on locally on certain parts of big data, the new methods are capable of utilizing large volumes of data fully to discover hidden patterns in complex biological systems.

We consider two types of machine learning approaches to reconstruct the network. The first type of approach is based on regression models, which aim to learn interaction between species from their linear regression relationship and construct the network framework using the estimated regression coefficients. The second type of approach is based on graphical models which estimate the precision matrix (the inverse of the covariance matrix) among all the species and infer the network structure based on the joint Gaussian assumption. Under this type of approach, we will further employ a variety of methods, evaluate their performance, and compare them with existing methods.

To further validate our analysis in practice, we apply all the new methods to a real study. The data set consists of time-series observations of 20 species abundances collected at Lake Constance in central Europe, which are population biomass (µgC/*m*^3^). We compare the reconstructed networks by machine learning algorithms to the known trophic network structure using four standard metrics (precision, recall, F1 score, network density) as defined in Section 3.3. In addition, we use the resampled data to generate highly-ranked interactions and evaluate their consistency across 7 methods and 5,000 learning processes. Based on our experiments, among all these methods under comparison, the best precision, recall, and F1 score were 0.48, 0.97, and 0.64, respectively, all achieved by the penalized regression methods. For instance, when the smoothly clipped absolute deviation (SCAD) used with AIC (Akaike Information Criterion), about 85% (SD = 0.0205) of the known interactions were captured in the reconstructed networks on average, and 60% of the retrieved interactions were consistently captured by all methods throughout 5,000 multiple learning processes. This study establishes the automated data-driven framework to reconstruct ecological networks while justifying the importance of choosing the proper type of data (e.g., panel data) to learn precise reconstructed networks.

## 2. Materials and methods

### 2.1. Data

In this research, we apply the machine learning methods to two different types of datasets from Lake Constance (Boit & Gaedke, 2014) for reconstructing ecological networks, validating the results, and evaluating the performance of algorithms. First, we use the abundance (µgC/*m*^3^) time-series data of 20 different groups of species in Lake Constance to reconstruct the ecological networks and capture biological interactions, which include both trophic and non-trophic interactions between species. The original panel data of species abundances are reduced to 1 year from 10 consecutive years (1987 - 1996) on a standardized time axis because, in this ecosystem, the inter-annual variability is much smaller than the seasonal variability (Boit & Gaedke, 2014). Second, an adjacency matrix of the trophic network (i.e., food web) is used to examine performance of models. The empirical dataset used for this study contains 20 functional guilds in Lake Constance and 365 measurements. In our analysis, we split the 365 observations into two parts: 75% are used to train and tune the models, and 25% are set aside as a validation set to test prediction accuracy.

### 2.2. Regression-based approach

Assume that there are *p* species in the ecological system. The observed values of all the species are represented together by a *p*-dimensional vector ***X*** = (*X*_1_,…, *X*_*p*_), where each species *X*_*i*_ contains *n* observations over time for species i and is treated as a random variable. We model the ecological networks between species by a graph *G* = (Γ, *E*), where Γ = {*X*_1_,…, *X*_*p*_} is the set of nodes and *E* is the set of edges in Γ × Γ. Our goal is to reconstruct the biological interaction networks in an ecological system, i.e., to identify possible connections among the nodes. We first consider a regression-based approach. For each species *X*_*i*_, we propose to learn its relations with other species by minimizing the penalized least squares,

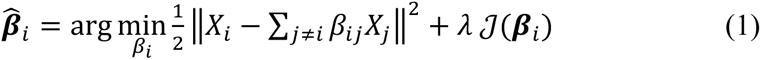

where the first term is the squared loss, the second *J*(***β***_*i*_) is a penalty function to control the complexity of the fitted model, and *λ* > 0 is the tuning parameter to balance the data fit and the model complexity. The penalty term plays a key role in high dimensional regression models to achieve high accuracy and interpretability. After solving (1) for all the species separately, we aggregate all the estimated regression coefficients 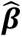 to reconstruct the networks. In particular, if 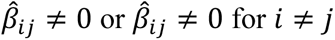, then there is an edge connecting species *i* and species *j*. Here we assume that the graphs are undirected.

The above learning algorithm is motivated by the neighborhood selection of Meinshausen and Bühlmann (2006), which estimates the structure of a undirected graph based on the LASSO regression relationship between the nodes. Here we consider a wider range of penalty functions to reconstruct biological interaction networks: the least absolute shrinkage and selection operator (LASSO) (Tibshirani, 1996), adaptive LASSO (Zou, 2006), ridge regression (Hoerl & Kennard, 1970), elastic net (Zou & Hastie, 2005), smoothly clipped absolute deviation (SCAD) (Fan & Li, 2001)), Gaussian graphical models (GGMs) (Dempster, 1972). See Figure 1 for the detailed workflow of the algorithms.

**Figure 1.**
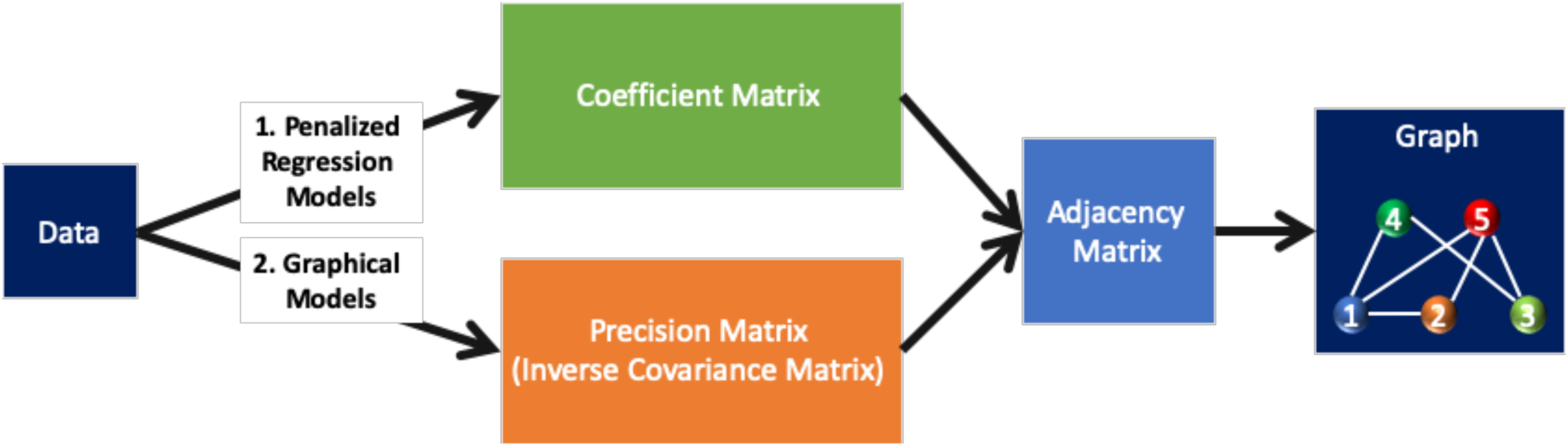
The diagram that describes the process of reconstruction algorithms using penalized regression and graphical models.

Different penalty methods have different properties. For example, the LASSO can improve the ordinary least squares (OLS) in terms of prediction accuracy and model interpretability by enforcing sparsity, but the estimators can be biased and lack consistency in variable selection, resulting in an overly dense graph with high false positives. Alternative penalized methods such as the elastic net (Zou & Hastie, 2005) and the SCAD (Fan & Li, 2001) can reduce estimation bias and have better theoretical properties such as variable selection consistency and oracle properties (Zou 2006). For optimal performance, the tuning parameter *λ* must be adaptively selected from the data. We will use Akaike Information Criterion (AIC; Akaike, 1974), Bayesian Information Criterion (BIC; Schwarz, 1978), and Cross validation (CV; Efron & Tibshirani, 1997; Hastie et al., 2013; Picard & Cook, 1984) to select the best *λ* in the penalized regression models. The details about the algorithms are summarized in Table 1.

**Table 1.**
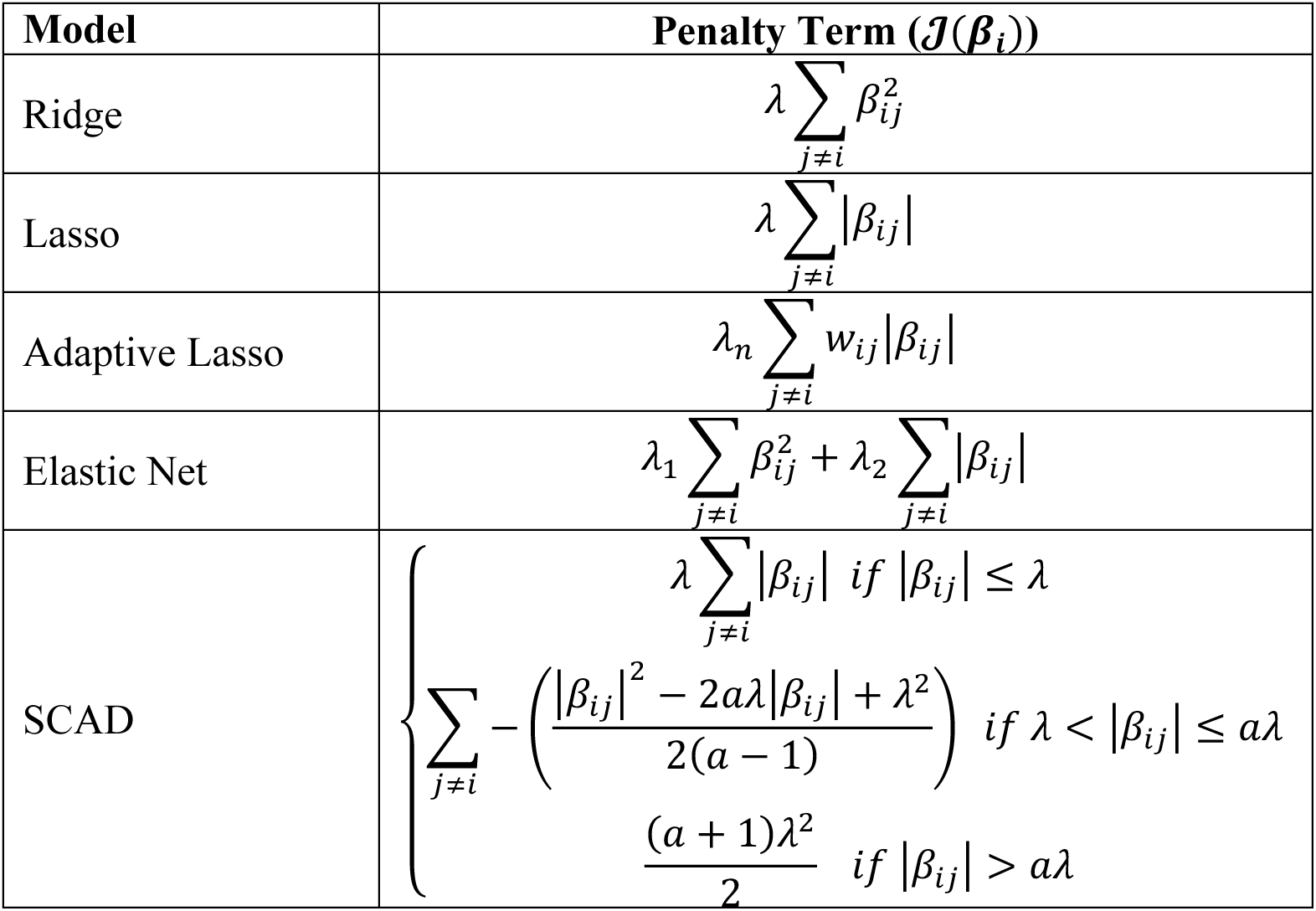
The penalized terms for different penalized regression models. Note: **λ** is a nonnegative regularization parameter, which controls sparsity, *W*_ij_ is a weight vector, and *a* is some number such that *a > 2*.

### 2.3 Graphical-Model based approach

The second approach is based on learning graphical models. Assume that ***X*** = *X*_1_,…, *X*_*p*_( have a multivariate distribution with mean **μ** and a positive definite covariance matrix Σ. If ***X*** is jointly distributed as the Gaussian, then a pair of nodes will have an edge if and only if they are conditionally dependent given the presence of the other nodes. In other words, If *X*_*i*_ and *X*_*j*_ are conditionally independent given all the other *X*’s, then entry *(i,j)* in the inverse covariance matrix Σ^−1^ will be zero (Lauritzen, 1996). If the Gaussian assumption does not hold, then having an edge between two nodes indicate a partially correlated relationship. Based on the above, we consider two graphical models – the GGM and the graphical lasso – to estimate the inverse covariance matrix and reconstruct the structure of the ecological network.

Let the inverse covariance matrix Θ = Σ^−1^, which is also known as the precision matrix. Let *S* be the sample or empirical covariance matrix, estimated from the data. Then, the GGM and graphical lasso maximize the following log-likelihood, except for the GGM, *ρ* = 0:

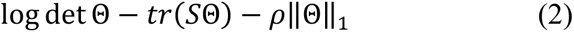

Here, det and tr denote the determinant and the trace of a matrix, respectively, ‖Θ‖_1_ is the *l*_1_ norm of a vector, and *ρ* > 0 is a tuning parameter to balance the tradeoff between the likelihood fit and the sparsity of the precision matrix. To select the best *ρ* in the graphical lasso, we use Akaike Information Criterion (AIC), Bayesian Information Criterion (BIC), and Cross validation (CV) to select the best *ρ* in the graphical lasso. There are multiple approaches to solve (2), and the R packages used in this study are summarized in Table 3.

## 3. Algorithms and implementation

In our analysis, we implement two ML approaches, the regression-based approach in Section 2.1 and the graphical model approach in Section 2.2, to capture species interactions and reconstruct ecological networks. For each approach, various penalized methods are implemented, and totally we end up with five penalized regression methods – the lasso, adaptive lasso, ridge regression, elastic net, and SCAD – and two graphical methods – GGMs and the graphical lasso (See S1 in SM for detailed descriptions of these methods).

Before fitting the model for each method, we need to first scale the panel data to count for the difference in magnitude of biomass across different groups of species. For the penalized regression methods, since the estimated are biased due to shrinkage effects, we refit the ordinary least squares (OLS) regression (See S1.1 for description of OLS refit) after variable selection to obtain unbiased estimated for reconstructing the ecological network.

### 3.1. Multiple learning processes

For the regression-based method, we randomly split the dataset into two parts, the training set for model fitting and coefficient estimation, and the test set for performance evaluation. To generate robust estimates of regression coefficients and reduce variability in results due to the random splitting, we repeat the process 5,000 times. For each process, we randomly sample 75% of the original data set to create a training set, which is used to fitting the penalized regression model and estimate the regression coefficients. To make the repeated results comparable among different methods, we set the same seed number for each random splitting, so that all the methods are trained with the same training set in each of the 5,000 processes.

For each training set, one species is regressed against all the rest of the species, and each species takes turns to be a dependent variable. Excluding the intercept coefficient, we obtain 19 coefficient estimates that correspond to 19 predictor species. Then, only using predictor species whose coefficient estimates are not 0, the OLS regression is refitted. When each of 20 guilds is finished taking turns, a 20-by-20 coefficient matrix can be formed, which has zero values for diagonal elements because no species was regressed against itself assuming there is no cannibalism in each group of species. Each learning process involves fitting a penalized regression model and the OLS refitting, so at the end of 5,000 repeated learnings for each method, 5,000 sets of a 20-by-20 coefficient matrix are obtained.

For the two graphical models, the entire data set is used to fit the model. Consequently, the multiple learning process will not change the results from graphical models. Fitting the graphical model generates three core elements needed to reconstruct ecological networks: covariance, inverse covariance, and partial correlation. The initial product of these graphical models is the precision matrix (also called the inverse covariance matrix), which can be inverted to produce the covariance matrix. Also, we use a ‘wi2net’ function in qgraph package in R statistical software (version 3.5.3) to obtain a partial correlation matrix from the precision matrix. For each of the seven ML methods, after computing all matrices, we take the absolute value of the resulting coefficient, covariance, precision, and partial correlation matrices and convert them into symmetric and binary matrices. To make reconstructed networks from both penalized models and graphical models comparable, all diagonal elements of the resulting matrices were set to zeros.

### 3.2. Reconstructing ecological networks

To reconstruct ecological networks, we use the coefficient matrices computed from the penalized regression models, and for graphical models, we use the covariance, inverse covariance, and partial correlation matrices. The sign and value of each element in these matrices and the asymmetricity of matrices can suggest interaction types, strengths, and directions of the interaction. However, in this study, we controlled for different interaction types (e.g., mutualism, commensalism, and antagonism), strengths, and directions. To investigate the existence of an interaction between two species in the reconstructed ecological networks, symmetric adjacency matrices obtained from penalized regression models and graphical models were used as the foundations to assemble the networks.

Each matrix can be visualized as an ecological network. Since we repeat the learning process 5,000 times, the results can be aggregated effectively by combining information drawn from 5,000 matrices for each method. Instead of arbitrarily choosing one matrix to represent the entire learning processes or taking the simple average of these matrices, we assemble the 5,000 matrices and retain interactions that are captured repeatedly with a high frequency, say, 99% of the time. By doing so, our algorithm is capable of summarizing the results from multiple processes and reconstructing ecological networks based on the *strongest* interactions among species. In addition, we label those interactions that were consistently identified by all the five penalized models and two graphical models at least 99% of the time as ‘highly-ranked’ and examine them furthermore.

### 3.3. Evaluating machine learning algorithms

To evaluate performance of the proposed machine learning methods for network reconstruction, we use the following metrics: precision, recall, F1 score, and network density. These performance metrics are calculated by counting true positives, false positives, and false negatives (Table 2). In particular, true positives are existing interactions that are captured by reconstructed networks. False positives are interactions that are falsely identified as existing in reconstructed networks, and they can be thoughts as “potential” interactions yet to be known. Similarly, false negatives are existing interactions that are missed by the method. Using these concepts, the precision is defined as the ratio of correctly identified interactions to all identified interactions in the reconstructed network, and the recall as the ratio of correctly identified interactions to all interactions in the empirical network. The F1 score is the harmonic mean of the precision and recall. The network density is the portion of connected links among all possible interactions in the reconstructed network.

**Table 2.** Summary of model performance metrics and their calculation methods.

| Performance metric | Calculation |
| --- | --- |
| Precision | Precision = (true positives)/(true positives+false positives) |
| Recall | Recall = (true positives)/(true positives+false negatives) |
| F1 score | F1 score = 2*(precision*recall)/(precision+recall) |

### 3.4. Software

We use the statistical software R (version 4.1.3) to implement machine learning methods. The names of R packages used in this study are summarized in Table 3. Our R code for data analysis, including data preprocessing, model fitting and tuning, and evaluation, are available upon request.

**Table 3.** Summary of machine learning algorithms used in this study and their respective package interface in R statistical software.

| Method |  | Reference | Package Interface |
| --- | --- | --- | --- |
| Penalized Regression | Ridge Regression | (Hoerl & Kennard, 1970) | MASS |
|  | Lasso | Tibshirani (1996) | lars |
|  | Adaptive Lasso | Zou (2006) | MASS, lars |
|  | Elastic Net | Zou and Hastie (2005) | glmnet |
|  | SCAD | Fan and Li (2001) | ncvreg |
| Graphical Models | GGM | Dempster (1972) | gRim |
|  | Graphical Lasso | Friedman, Hastie, and Tibshirani (2008) | glasso |

## 4. Results

### 4.1. Prediction accuracy of the penalized regression model

The five penalized regression models - the lasso, adaptive lasso, ridge regression, elastic net, and SCAD – were implemented to predict abundances of 20 different groups of species by regressing one species against all the other species for all 20 guilds. Each species took turns to be the response variable in a regression model, so we obtained 20 different regression models. Since we repeated the learning processes 5,000 in order to get robust estimates of the regression coefficients, the predicted values of each regression model were averaged and plotted with the observed values on the same plot to illustrate the prediction accuracy of our penalized regression models (Fig. 2).

**Figure 2.**
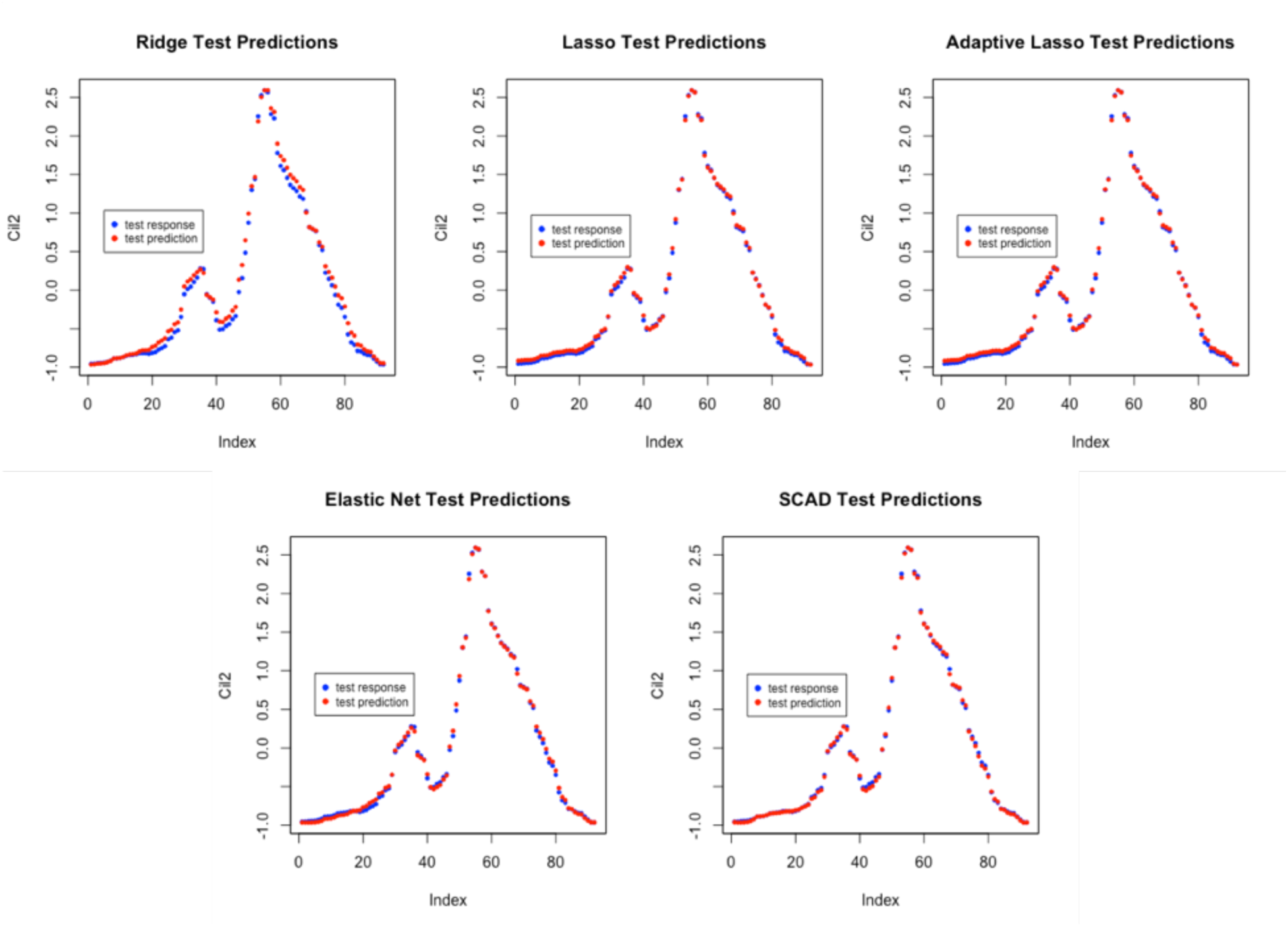
Predictions of abundances of Cil2 species using the 5 penalized regression models.

The prediction accuracy is measured by the mean squared error (MSE), which is the square of mean differences between the observed response values and the predicted response values on the test set. As shown in Fig. 3, the accuracy results from the five penalized regression methods were quite close to each other: the smallest mean MSE of 5,000 learning processes was 0.36 and the largest mean MSE was 0.047, corresponding to different selection criteria (CV, AIC, BIC) for parameter tuning. Overall speaking, the SCAD with CV tuning works best for prediction with the lowest MSE 0.036.

**Figure 3.**
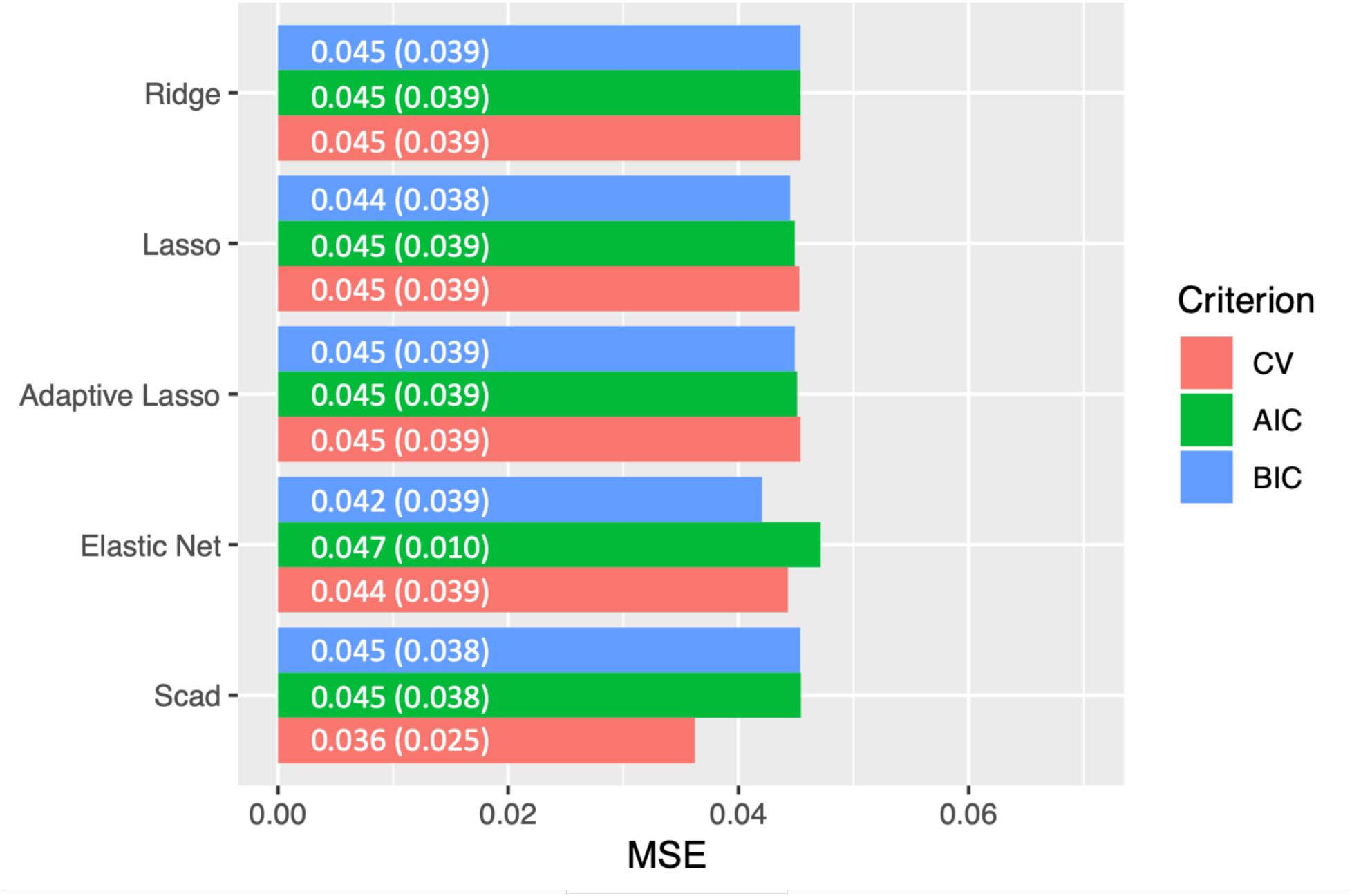
The mean MSE and the standard deviations of 5,000 learning processes for 5 penalized models by different tuning criteria.

### 4.2. Performance on network reconstruction

Table 4 summarizes the network construction performance for three penalized regression models and one graphical model, in terms of precision, recall, and F1 scores (see ST2. for full results). The network density was calculated to compare densities of reconstructed networks to that of the original network. The highest score of precision, recall, and F1 score across all tuning criteria were 0.48, 0.97, and 0.64, respectively. Overall speaking, the LASSO gives the best performance among the methods in terms of precision, recall, and F1 scores, while the elastic net produces the sparsest graph. For all the three penalized regression models, CV overall led to better performance than AIC and BIC, and the latter two give similar performance.

**Table 4.**
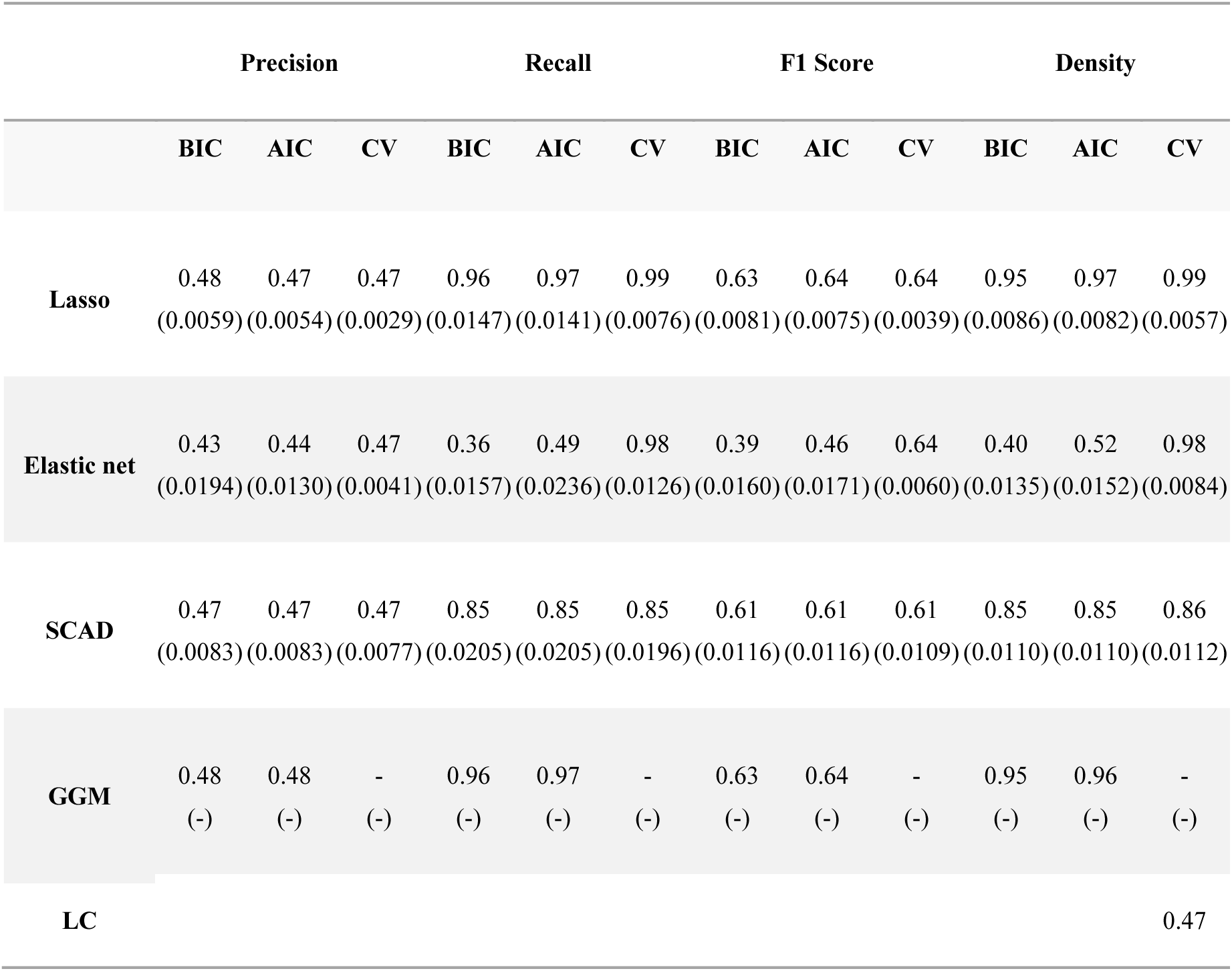
The mean and standard deviation of precision, recall, F1 score, and density from 5,000 learning processes for the lasso, elastic net, SCAD, and GGM.

The density of Lake Constance food web was measured after it was transformed into undirected network. Compared to the empirical network density 0.47, the overall densities of reconstructed networks were higher for all the methods except that of the elastic net with BIC (which was 0.40). However, it is important to note that reconstructed networks are different from the empirical food web because they capture both trophic and non-trophic interactions between species, which may lead to higher network densities.

### 4.3. Reconstructed networks

In Figure 4, we visualize the reconstructed networks from the elastic net based on BIC and AIC, and compare them with the known Lake Constance food web. In each network, the nodes represent 20 different groups of species, and the edges represent interactions between species. For visualization, the trophic levels increase counter-clockwise from Alg1 (Single-cell algae) to Lep (Cladocerans), and the node colors are differentiated by seven different functional groups (For more details about these functional groups, see Boit et al. 2012).

**Figure 4.**
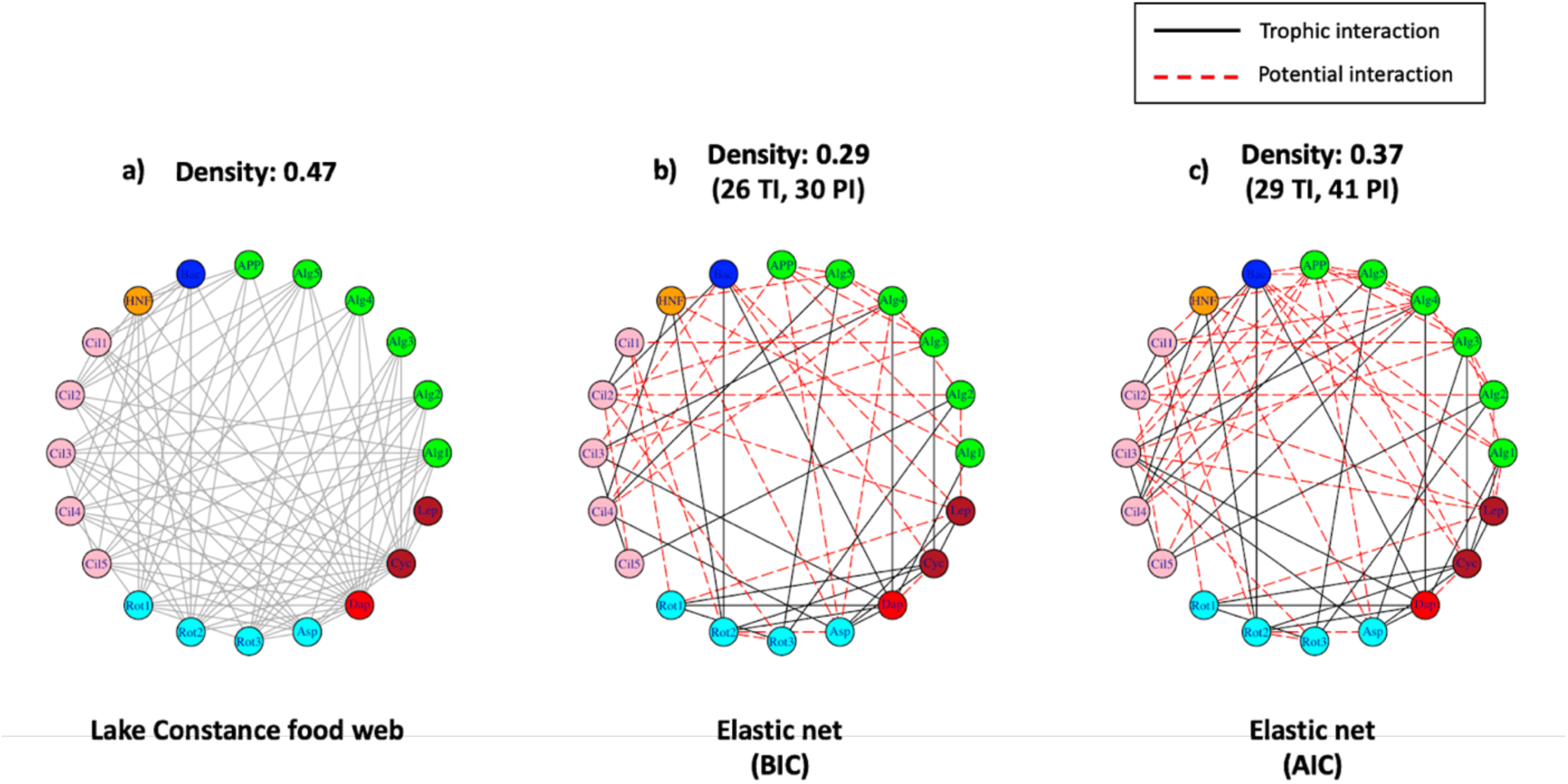
The reconstructed networks made by penalized regression and graphical methods along with empirical food web. The realizations of the reconstructed networks are based on the interactions that were captured 99% of the 5,000 learning processes. In that sense, the density values for the reconstructed networks were lower than those from Table 4 because interactions that were not captured 99% of the time were excluded in these network visualizations.

Figure 4a shows the empirical Lake Constance food web (Fig 4a), where each edge describes a trophic interaction between species. Figures 4b and 4c show the reconstructed networks, with black solid edges and red dotted edges representing trophic interactions and potential interactions, respectively. Each reconstructed network identifies those interactions captured 99% out of 5,000 matrices from the multiple learning.

Although the empirical food web consists of trophic interactions only, two reconstructed networks include more than feeding relationships (e.g., non-trophic interactions) represented by potential interactions (dotted red lines). Considering the incompleteness of the reference trophic network, these newly found interactions are likely to be either unknown trophic interactions or known/unknown non-trophic interactions. The most obvious examples of non-trophic relationships shown in reconstructed networks can be found in interactions among primary producers (i.e., APP and Alg1-5), which do not exist in the empirical food web. In Figure 4, the density of the reconstructed network from the elastic net based on AIC (Fig. 4c) is found to be, which is closer to that of Lake Constance food web than 0.29 based on BIC (Fig. 4b). Moreover, two reconstructed networks are comprised of more potential interactions, which are not known trophic interactions, than known trophic interactions. In addition, from Figure 8, we observe that reconstructed networks using elastic net showed variations in performance metrics and densities across different tuning criteria (BIC, AIC, and CV). For example, the densities of reconstructed networks using elastic net with BIC, AIC, and CV were 0.29, 0.37, and 0.91, respectively.

### 4.4. Identifying highly-ranked signals

Trophic interactions include interactions identified in the reconstructed networks, and also those potential interactions not discovered yet, which are not shown in the empirical network but detected by the reconstructed network.

To assess the consistency in results across all the methods over multiple learning processes, we identify highly-ranked interactions, which are quantified by computing the number of times of any interaction captured by each method and summarizing the frequencies of interactions captured by 99% of the time by the number of methods (Figures 5-7). Since the densities of reconstructed networks based on the BIC were the sparsest, the methods with BIC tend to have the fewest number of interactions captured 99% of the time by all the methods. In contrast, those based on CV were the least sparse, and consequently, they have the largest number of interactions captured 99% of the time by all the methods.

**Figure 5.**
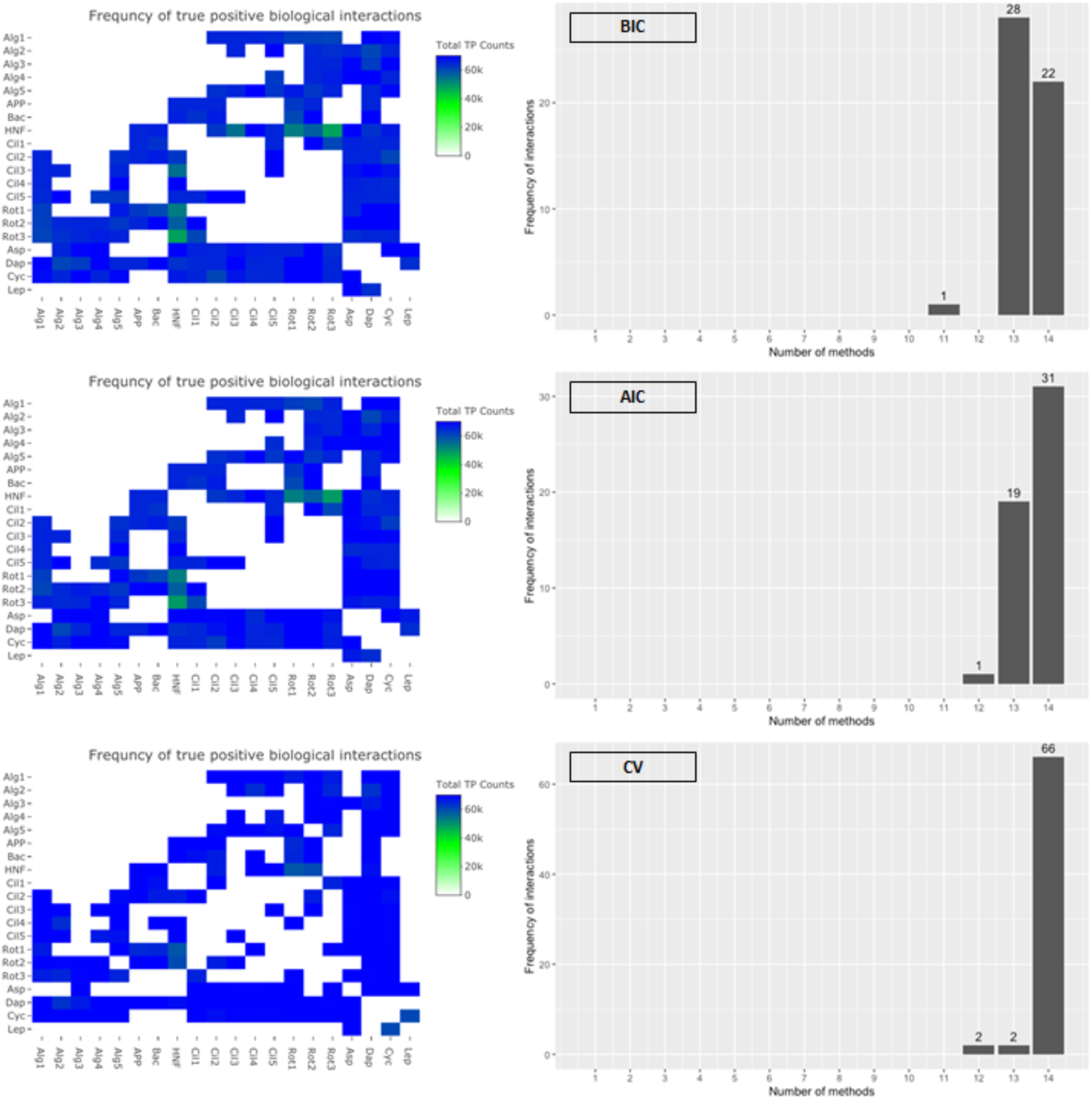
The heatmap of true positive (trophic interaction) values of the reconstructed networks from the penalized regression models and the graphical models (left). The bar graph shows the frequencies of true positives captured by 99% of the time by all approaches of those penalized regression models and the graphical models (right).

**Figure 6.**
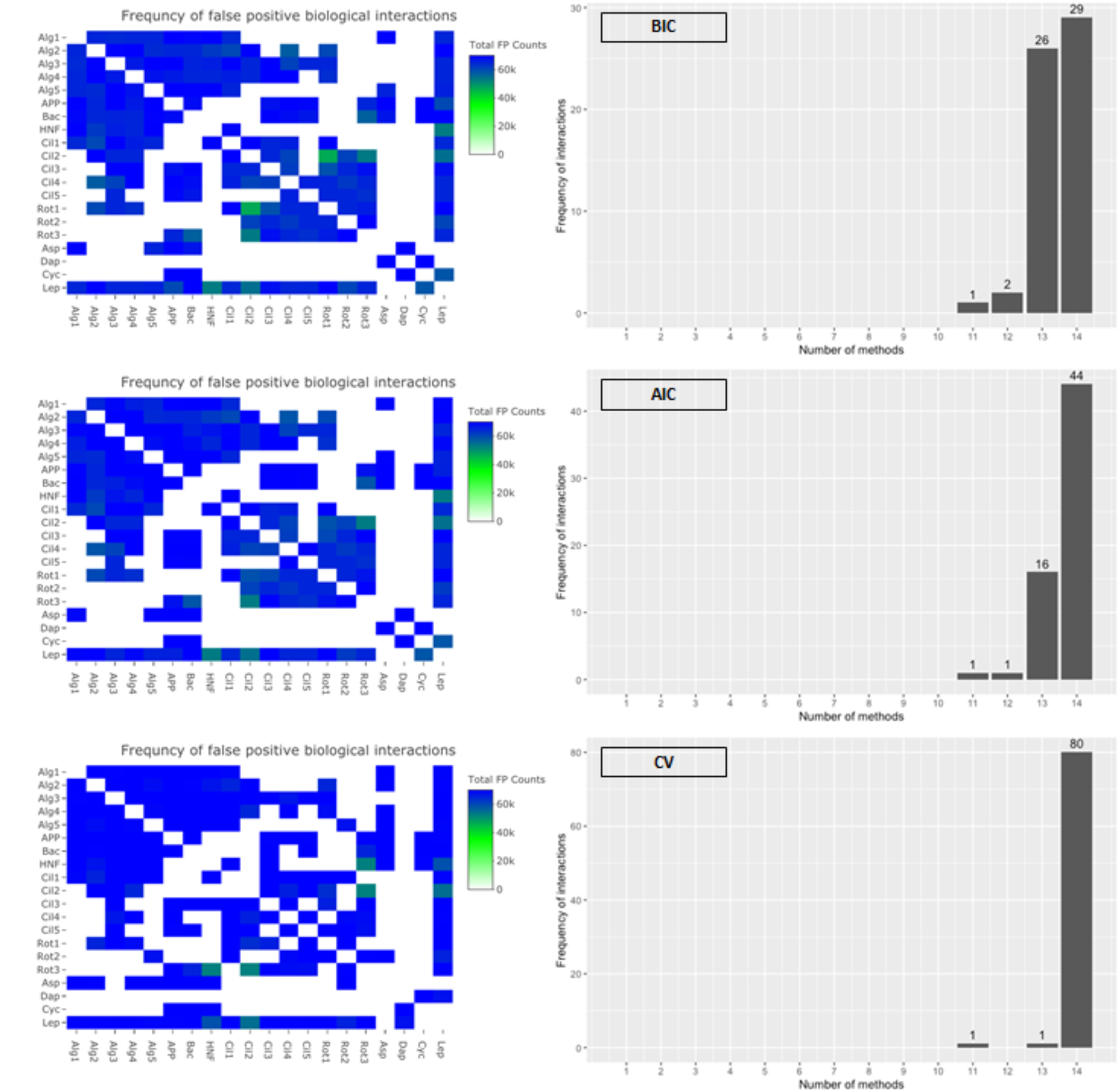
The heatmap of false positive (potential interaction) values of the reconstructed networks from the penalized regression models and the graphical models (left). The bar graph shows the frequencies of true positives captured by 99% of the time by all approaches of those penalized regression models and the graphical models (right).

**Figure 7.**
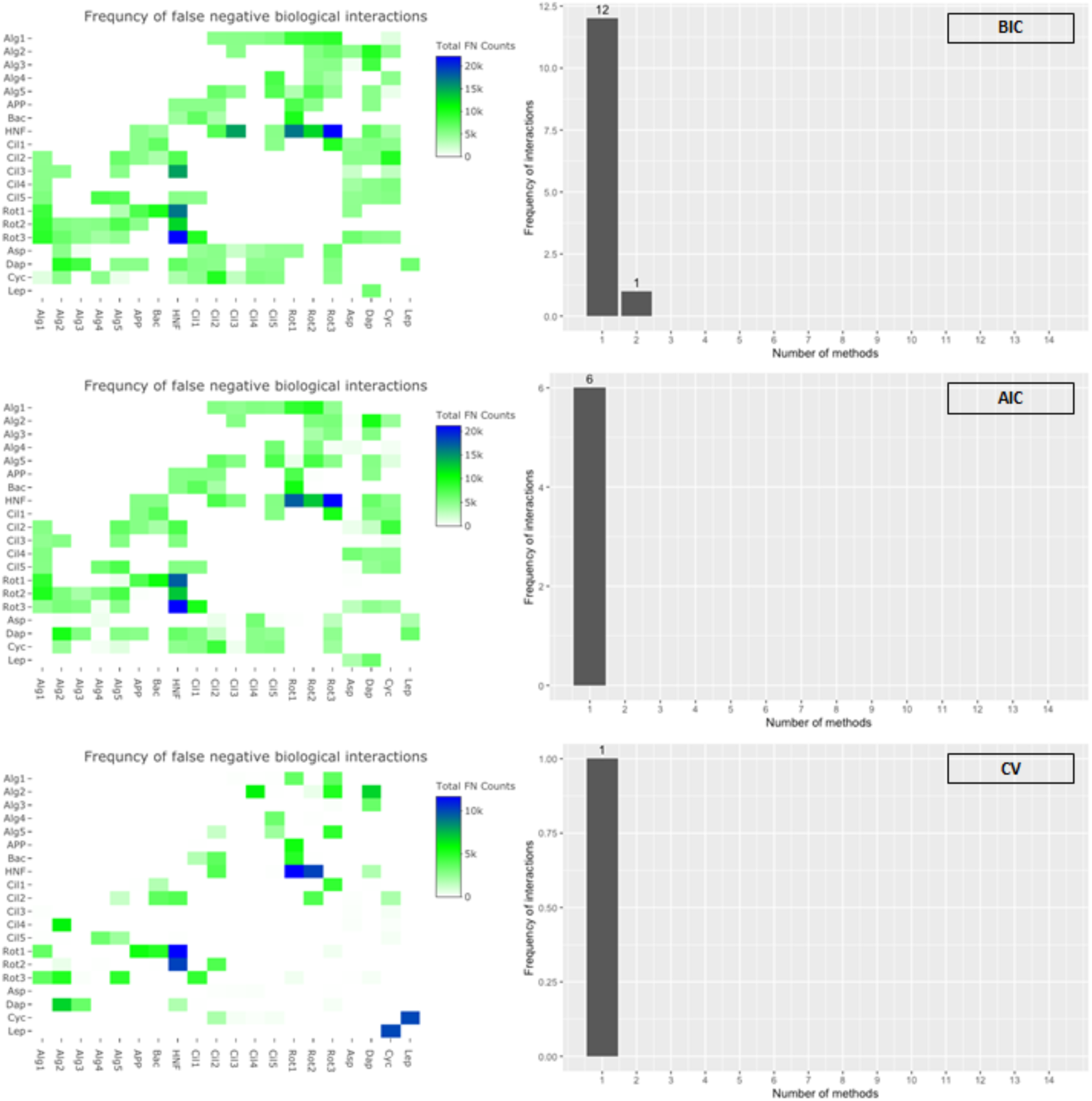
The heatmap of false negative values of the reconstructed networks from the penalized regression models and the graphical models (left). The bar graph shows the frequencies of true positives captured by 99% of the time by all approaches of those penalized regression models and the graphical models (right).

Out of 190 possible interactions from the symmetrized Lake Constance food web of 20 groups of species, 90 links are known trophic interactions and 100 links can be considered ‘potentially new’ interactions. Among 90 known trophic links, 22 (≅24% of all existing trophic interactions) trophic interactions were captured 99% of the time by all the methods based on the BIC, 31 (≅34%) based on the AIC, and 66 (≅73%) based on the CV (See histograms in Fig. 5). For potential interactions, 29 (29% of all potential interactions) interactions were captured 99% of the time by all methods based on the BIC, 44 (44%) based on the AIC, and 80 (80%) based on the CV. Lastly, for the false negative interactions, 12 (≅13% of all existing trophic interactions) and 1 (≅1%) interactions are captured 99% of the time by one and two methods respectively based on the BIC, 6 (≅6%) interactions by only 1 method based on the AIC, and 1 (≅1%) interaction by 1 method based on the CV.

## 5. Discussion

Machine learning algorithms have been applied in ecology over decades, but with somewhat limited purposes. Especially at the community scale, these techniques have not shown promising results for network inferences (Faisal et al., 2010; Sander et al., 2017) and making predictions with ecological data (Shan et al., 2006) because of their overall relatively low performance. This work shows that, using species abundance time-series data of 20 different groups of species in Lake Constance, the reconstructed ecological networks based on five statistical penalized regression methods and two graphical methods suggest the promise of machine learning algorithms for studying ecological data and discovering underlying biological interactions among species. Although for some methods, precision, recall, and F1 score of the reconstructed network were relatively low overall compared to other methods used in this study, the overall performances were higher than the previous comparative study (Faisal et al., 2010; Sander et al., 2017) of different ecosystems. Also, most reconstructed networks not only captured known trophic interactions among species but might have discovered unrevealed interactions among some species, which may lead to further research to study new relations among those species.

### 5.1. Advantages of using species abundance panel data

The presence-absence data have been collected and used to show patterns of species distribution by accumulating the data regarding whether each individual of species exists or not on a certain site at the time of data collection. Although it can be useful to collect presence-absence data when the study budget is limited (Joseph et al., 2006), the information given by the presence-absence data can be limited that might cause relatively less precise network reconstruction. On the other hand, the panel data of species abundances have more information by providing multiple observations on abundances of each group of species over time in the shape of gradients. Although most of the previous studies that have inferred ecological networks used presence-absence data which is binary data (Faisal et al., 2010; Sander et al., 2017), more insights about species interactions to better reconstruct community networks can be drawn if time-series abundance data of groups of species which is panel data were used instead.

There has been a study to clarify how these different types of data can affect habitat preference evaluation (Fukuda et al., 2012) which showed better performances based on species abundance data than binary data. However, regardless of its importance, no research has examined the effects of the types of data on performance of reconstructing ecological networks based on machine learning methodologies. Hence, it is crucial to shed light on the unanswered question of whether a certain type of data can be more beneficial. Although further studies with various types of data from a specific ecosystem are required to be tested for reaching a comprehensive conclusion, this study contributes to existing literature to resolve this challenge by showing enhanced performances in general.

### 5.2. Prediction accuracy of the penalized regression model

The penalized regression models are widely used to make predictions especially for high-dimensional data or for highly correlated predictors because these penalized regression methods are effective in reducing variance of the fitted models (Tibshirani, 1996) and avoid the overfitting problems of the ordinary least squares regression. Depending on the choice of the penalty function, the predictive power of a model can be significantly improved (Fan & Li, 2001), and a parsimonious model can be produced and offers better interpretability about the underlying mechanism of data generation (Zou & Hastie, 2005). We have shown that the penalized regression models of interest can capture the dynamics of ecological systems and have high prediction accuracy (Fig. 2) with time-series data of species abundance.

For all 5,000 learning processes, the overall prediction accuracies (i.e., 1-MSE) were high on average ranging from 0.95 to 0.96 (Fig. 3). The overall high prediction accuracies were apparent in all penalized regression methods of our interest. In specific, the high prediction accuracy of the penalized regression models using 1-year of aggregated panel data from Lake Constance suggests that the coefficient estimates of these regression models are highly accurate to represent the empirical data. It also implies that reconstructed ecological networks based on these coefficient estimates can precisely represent the species interactions in the empirical data, supporting the fact that the reduced 1-year data from the original 10-year data is sufficient. It was also known that interannual variability was much smaller than the seasonal variability within the original data (Boit & Gaedke, 2014), and the high prediction accuracy suggests that the selected penalized regression models well captured the seasonal variability.

### 5.3. Performance of the machine learning (ML) models

The reconstructed networks derived from the machine learning (ML) models include both trophic and non-trophic interactions because the ML models were used to detect the existence of species interaction between a pair of species. More specifically, the penalized regression models capture the relations among species based on the coefficient values if the coefficient estimates are not zero, and unless the graphical models estimate that the conditional independence does not exist between any two species, the two species are assumed to have a relation whether it is trophic or non-trophic.

Our model being able to capture both trophic and non-trophic interactions is supported by the density values of the reconstructed networks from the different models. Most of the density values are higher than the density of the reference network, which solely describes the trophic relationships among species. If the fully investigated interaction network of Lake Constance were available as a reference network, our results would have been much better in terms of the performance metrics. In other words, the three performance metrics are highly likely to increase if the reconstructed networks were compared to the comprehensive interaction network, since it will be more probable for potential interactions to be identified as existing interactions, it will increase the number of true positives while decreasing the number of false positives and false negatives. Then, the precision, recall, and F1 score values will increase accordingly.

Although the empirical network only includes trophic interactions, comparing reconstructed networks to the empirical network is still relevant. A high recall value can suggest that the reconstructed model can capture existing trophic interactions very well, and a moderate precision value can imply existence of potential non-trophic species interactions. If the network density is higher than that of the empirical network for a reconstructed network with a moderate precision value and high recall value, then it confirms that the respective model is able to capture trophic and non-trophic interactions. Therefore, it is still worth examining these performance metrics by comparing each reconstructed ecological network to the empirical network. Again, if a more complete biological interaction web were used as a true reference, then model performances might be better except for the fact that a complete biological interaction network especially in a system with a large number of species is rarely assembled.

### 5.4. Highly-ranked signals

When it comes to applying ML methods to reconstruct ecological networks, the consistency of network structures needs to be examined across different methods over ample iteration. Highly-ranked signals can be measures of the consistency to verify whether captured interactions are strong enough signals agreed among many models repeatedly. Because the study is data-driven including many unknown interactions, even relatively weaker species interactions that were captured by our models pose some questions that can be explored further in future studies.

Since the network densities of most methods found to be 0.9 or close to 0.9 indicates that all species have interactions with other species. When we used a threshold value to remove weak interactions, the density values became significantly lower for the graphical lasso and the GGM. However, it is hard to justify the reason for arbitrarily setting a threshold value. Furthermore, since our aim is to capture interactions regardless of their types, it makes more sense that all species are linked in the reconstructed networks.

### 5.5. Effect of three different tuning criteria (BIC, AIC, and CV)

When the elastic net with BIC was applied to reconstruct the ecological network, it tends to capture strong signals rather than weak signals. It helps to explain why this network has the lowest density. However, when CV is used as a criterion, the algorithm captured both strong and weak signals (Fig. 8).

**Figure 8.**
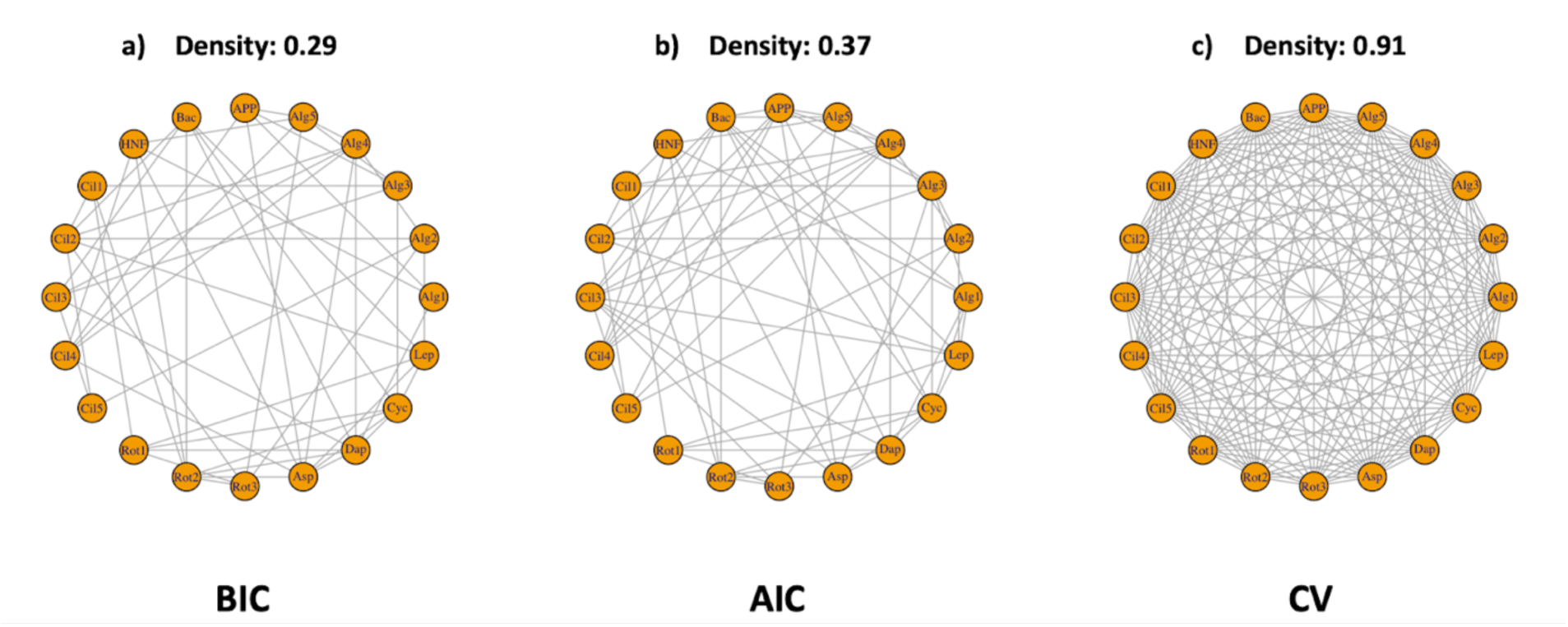
Network realizations of the elastic net by different tuning criteria.

The precision, recall, F1 score, and density were compared using the BIC, AIC, and CV for choosing the tuning parameter to control for the amount of the shrinkage (Table 3). For the penalized regression models, all three criteria are used to select the tuning parameter, but for the graphical models, only the AIC and BIC are used for tuning. For all models, tuning parameters that minimize the AIC, BIC, or the CV values were chosen. Because the tuning parameter controls the density of the model, chosen parameters will directly affect the density of the reconstructed networks. Even for the same machine learning algorithms, the density values will differ depending on the criteria to control the tuning.

In general, a model chosen by minimizing the BIC gives the sparsest result compared to models chosen by minimizing the AIC or the CV errors. For example, the densities of the reconstructed network from the elastic net after the refitting are 0.29, 0.37, and 0.91 based on the BIC, AIC, and the CV, respectively. The same pattern was prevalent for many other machine learning methods. The general pattern is also clear from the heatmaps in Figures 5-7.

The density ranges from 0.40 to 0.99, and the elastic net had the density of 0.36 and 0.47 based on the BIC and AIC, respectively. Most density values were around 0.9, so we explored the density values by looking through a sequence of tuning values to see if we could achieve a density around 0.40. Although it is possible to achieve a density of 0.40, the performances of models based on the precision, recall, and F1 score become worse.

## 6. Conclusion

Our network reconstruction algorithms demonstrate effective and better performance in network reconstruction than classic statistical method (i.e., Pearson’s correlation) and as well as others in existing literatures based on various types of empirical data (e.g., presence-absence data, count data, trait data, etc.) from different ecosystems. These new machine learning methods not only provide automated data-driven tools to reconstruct ecological networks, also prove importance of choosing the proper type of data (e.g., panel data) to obtain precisely reconstructed networks by using our algorithms. Furthermore, the ML algorithms are flexible in handling both directed and/or weighted networks, they have broad applications to further studies.

## AUTHOR CONTRIBUTIONS

H.K. conceived and led the project; H.K. collected the data (from existing sources); H.K., A.A.K., and H.H.Z. designed methodology; H.H.Z. offered methodological guidance throughout the project. H.K. and A.A.K. coded, designed, and performed computational experiments and simulations; H.K. illustrated the results; All authors analyzed the results; H.K. led the writing of the manuscript; All authors contributed critically to the drafts and gave final approval for publication.

## ACKNOWLEDGEMENTS

We would like to thank the Department of Ecology and Evolutionary Biology (EEB) and the Department of Mathematics at the University of Arizona. HPC infrastructure support was provided by the University of Arizona’s High-Performance Computing (HPC) system. This project has been supported by funding from U.S. National Science Foundation (TRIPODS 1839307).

## DATA AVAILABILITY STATEMENT

Data available from:

Boit, A., & Gaedke, U. (2014). Benchmarking Successional Progress in a Quantitative Food Web. *PLOS ONE*, *9*(2), e90404. https://doi.org/10.1371/journal.pone.0090404

---

**Table X.**
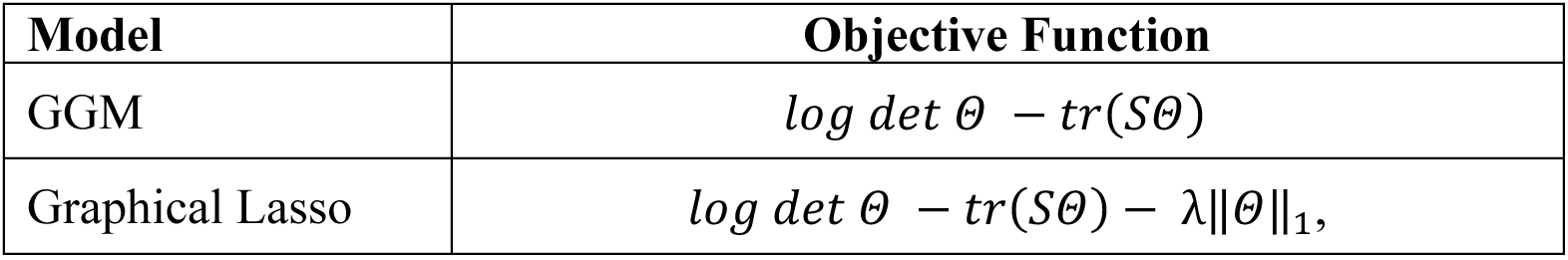
The likelihood functions for graphical models. Note: *Θ* is the precision matrix, *S* is the sample covariance matrix, λ is a tuning parameter that controls sparsity, *W*_*ij*_ is a weight vector, and *a* is some number such that *a > 2*.

